# Dynamics of giant vesicle assembly from thin lipid films

**DOI:** 10.1101/2023.07.02.547429

**Authors:** Joseph Pazzi, Anand Bala Subramaniam

**Affiliations:** Department of Bioengineering, University of California, Merced, Merced, CA 95343, United States

## Abstract

Giant unilamellar vesicles (GUVs) are micrometer-scale lipid assemblies that emulate key characteristics of biological cell membranes. GUVs can be obtained when solid-supported thin films of lipids are hydrated in aqueous solutions. However, a comprehensive understanding of their assembly dynamics has been lacking, impeding mechanistic insights. Here, we report the time dependence of the distribution of sizes and molar yield of GUVs obtained through a novel ‘stopped-time’ technique. We compare three commonly used techniques, PAPYRUS (Paper-Abetted amPhiphile hYdRation in aqUeous Solutions) gentle hydration, and electroformation. We demonstrate that all three techniques show sigmoidal yield curves. Yields increase monotonically before reaching a plateau, with surprisingly high yields 60 seconds after hydration. Gentle hydration shows limited time evolution in contrast to PAPYRUS and electroformation. Exploration of bud dynamics on the surfaces uncovers bud emergence, diameter growth, and merging phenomena. To provide a comprehensive explanation of our observations, we employ the thermodynamic budding and merging model. This work expands our understanding of GUV assembly dynamics and offers fundamental insights into the underlying thermodynamic principles governing this process.

## Introduction

Thin lipid films supported on solid surfaces form spherical buds > 1 µm when hydrated in low salt aqueous solutions. Once detached from the film, these buds close to form giant unilamellar vesicles (GUVs). Due to their comparable size to biological cells, ability to mimic the chemical and physical properties of plasma membranes, and ability to compartmentalize soluble molecules in their lumens, GUVs have found wide use in studies of membrane biophysics,^1–3^ synthetic biology,^4–6^ and in biomedical applications.^7–9^ Techniques that employ the hydration of thin lipid films^10, 11^ include gentle hydration or natural swelling which uses glass surfaces,^12, 13^ electroformation or electroswelling which uses conductive surfaces,^14–16^ gel-assisted hydration which uses surfaces coated with partially soluble polymers,^17–19^ and PAPYRUS, Paper-Abetted amPhiphile hYdRation in aqUeous Solutions, which uses nanocellulose paper.^20^ Of these techniques, only electroformation involves the active input of energy via the application of electric fields. ^14–16^ All the other techniques allow the thin lipid films to evolve under quiescent conditions after hydration.

We recently reported an experimental framework to quantify the distribution of sizes and molar yields of populations of GUVs using confocal microscopy and large data set image analysis.^20^ By standardizing experimental conditions and through statistical analysis, we showed that the yield of GUVs obtained using PAPYRUS was significantly higher compared to electroformation and gentle hydration on glass surfaces. We showed that the free energy change for forming spherical buds from bilayers wrapped around the nanoscale cylindrical fibers of nanocellulose paper is significantly lower compared to the free energy change for forming spherical buds from bilayers supported on flat surfaces. The low free energy cost explains the high yield.^20^ The method of quantification additionally allowed the measurement of the relative contributions and the elucidation of the mechanism of action of six assisting compounds on the yields of GUVs in salty solutions.^19^ Here, we use similar principles to develop a novel ‘stopped-time’ technique to study the time dependence of the evolution of the distribution of diameters and molar yields of GUVs. Time evolution data is critical to understand the mechanisms involved in the assembly of GUVs from thin lipid films.

We studied three different thin film hydration techniques, PAPYRUS, gentle hydration on flat borosilicate glass slides, and electroformation on indium tin oxide (ITO)-coated glass slides. To the best of our knowledge, this paper presents the first report of the time evolution of statistically representative populations of GUVs obtained from multiple thin film hydration techniques. Previous studies have focused predominantly on the electroformation technique and on investigating the time evolution of single to few buds on the lipid films.^16, 21–25^

We find that all three thin film hydration techniques produce an asymmetric distribution of GUV diameters for all time points. Large numbers of GUVs were present after the relatively short time of 1 minute. Over the course of 120 minutes, the distributions broaden due to the increase in the number of GUVs with diameters > 10 µm. The histogram of diameters could not be fit adequately by common functions used to describe the distribution of dispersed particles. We find that the number of GUVs with diameters < 10 µm showed non-monotonic behavior with time. The number of GUVs with diameters > 10 µm increased monotonically and reached a plateau by 120 minutes for PAPYRUS and electroformation. Gentle hydration showed limited increase in the number of GUVs with diameters > 10 µm. GUVs with diameters < 10 µm are ∼ 13 to 32 times more abundant than GUVs > 10 µm in diameter, with PAPYRUS showing the largest fraction of GUVs with diameters > 10 µm and gentle hydration showing the smallest fraction.

The molar yield of GUVs increased monotonically before reaching a plateau, demonstrating a characteristic sigmoid shape for all three techniques. The yield of GUVs at the plateau was 31 %, 16 %, and 22 % for PAPYRUS, gentle hydration, and electroformation respectively. The incorporation of new lipids into the pool of buds that form GUVs, that is the period of increase in yield, ceases within 30 and 60 minutes for PAPYRUS and electroformation respectively. For PAPYRUS and electroformation, coarsening of the buds to form GUVs of large diameters continues to occur up to 120 minutes after hydration. Increasing incubation time beyond 120 minutes results in a decrease in yield. Gentle hydration on glass showed both limited evolution in yield and limited coarsening compared to PAPYRUS and electroformation.

To understand the time evolution of the GUVs, we took time-lapse confocal microscopy images of the buds evolving on the surfaces. At the typical lipid surface concentrations that we used, we observe predominantly merging of the close-packed micrometer scale buds. The rate of merging decreased with time for all three surfaces, albeit merging of buds is minimal for gentle hydration on glass slides. We also prepared surfaces with a low concentration of lipids. On these surfaces, the low density of buds allowed us to clearly observe the emergence of isolated micrometer scale buds from regions of lipid films that were initially devoid of buds. Buds also slowly increase in diameter without any micrometer-scale merging. Nevertheless, even at these low concentrations, adjacent buds often merge with each other as they increase in diameter. A combination of the complex dynamics of bud emergence, diameter increase, and merging likely gives rise to the non-trivial shape of the distribution of diameters.

Our results show previously unappreciated commonality between the three thin film hydration techniques. We show that the data is consistent with the budding and merging model for the assembly of GUVs.

## Methods

### Materials

We purchased premium plain glass microscope slides (75 mm × 25 mm), glass coverslips (Gold Seal™, 22 mm × 22 mm) and large Petri dishes (Falcon™ Bacteriological Petri Dishes with Lid, 150 mm diameter, 15 mm height) from Thermo Fisher Scientific (Waltham, MA). We purchased indium tin oxide (ITO) coated-glass slides with dimensions of 25 mm × 25 mm and surface resistivity of 8-12 Ω/sq from Sigma-Aldrich (St. Louis, MO). Artist grade tracing paper (Jack Richeson & Co., Inc.) and hole punch cutters (Amon Tech) were purchased from Amazon Inc. (Seattle, WA).

### Chemicals

We purchased sucrose (BioXtra grade, purity ≥ 99.5%), glucose (BioXtra grade, purity ≥ 99.5%), and casein from bovine milk (BioReagent grade) from Sigma-Aldrich (St. Louis, MO). We purchased chloroform (ACS grade, purity ≥ 99.8%, with 0.75% ethanol as preservative) and poly(dimethyl)siloxane (Krayden Dow Sylgard 184 Silicone Elastomer Kit.) from ThermoFisher Scientific (Waltham, MA). We obtain ultrapure water (18.2 MΩ) from an ELGA Pure-lab Ultra water purification system (Woodridge, IL). We purchased 1,2-dioleoyl-*sn*-glycero-3-phosphocholine (18:1 (Δ9-cis) PC (DOPC)) and 23-(dipyrrometheneboron difluoride)-24-norcholesterol (TopFluor-Chol) from Avanti Polar Lipids, Inc. (Alabaster, AL).

### Lipid composition and concentration

The composition of the lipid mixture we used was DOPC:TopFluor-Chol at 99.5:0.5 mol %. We deposit 10 µL of a 1 mg/mL concentration of the lipid mixture (10 µg lipid, 17 nmol/cm^2^) onto a 9.5 mm diameter circular area of the substrates for typical experiments. We deposit 10 µL of a 0.25 mg/mL concentration of the lipid mixture (2.5 µg lipid, 4.25 nmol/cm^2^) onto a 9.5 mm diameter circular area of the substrates for the sparse buds experiments.

### Stopped-time technique

We followed a previously reported protocol to clean the substrates and assemble GUVs.^20^ We arrested the evolution of buds by harvesting the GUV buds at 1 minute, 10 minutes, 30 minutes, 60 minutes, and 120 minutes. We carefully disassembled the chamber by removing the top slide for gentle hydration and electroformation. We aspirated and expelled 100 µL of the hydrating solution 6 times on different regions covering the whole substrate. After the sixth aspiration, we collected all of the liquid ∼ 150 µL containing the GUVs and transferred the liquid into an Eppendorf tube. We immediately moved a 2 µL aliquot of the GUV solution into a chamber for imaging. The imaging chambers were constructed by covalently bonding custom-made square PDMS gaskets with dimensions of 6 × 6 × 1 mm (width × length × height) to glass microscope slides. To prevent the rupture of the GUVs on the glass, we passivated the surface using a solution of 1 mg/mL casein in 1× PBS buffer for 1 hour, and then washed away the unbound casein with 3 washes using ultrapure water. We placed 58 µL of a 100 mM solution of glucose and then 2 µL of the suspension of harvested GUVs into the passivated chamber. We sealed the chamber using a glass coverslip and allowed the GUVs to sediment for 3 hours before imaging. We used an upright confocal laser-scanning microscope (LSM 880, Axio Imager.Z2m, Zeiss, Germany) to collect images above the surface of the glass where the GUVs were sedimented. We excited the TopFluor® lipid dye contained in the membrane of the GUVs with a 488 nm argon laser. The fluorescence was collected using a 10× Plan-Apochromat objective with a numerical aperture of 0.45. We collected 64 images covering the entire area of the chamber using an automated tile scan routine with each image covering an area of 850.19 µm × 850.19 µm (3212 pixels × 3212 pixels). The routine used an autofocus feature to focus on the vesicles resting above the surface of the glass at each tile location. The confocal pinhole was set to 12.66 Airy Units which gave a confocal slice thickness of 80 µm. We conducted three independent repeats per time point for a total of 18 independent samples for each of the three substrates.

### Image processing and data analysis

We used a custom routine in MATLAB (Mathworks Inc., Natick, MA) to analyze the vesicles from the confocal tilescan images. We applied a threshold and a watershed algorithm to segment the fluorescent objects from the background. We obtained the equivalent diameters and the mean intensities of each of the segmented objects using the native *regionprops* function. To select GUVs from non-GUV lipid structures (multilamellar vesicles, nanotubes, aggregates), we analyzed the segmented objects based on the coefficient of variance of their intensity values. We defined objects that fell within 1 – 2 times the full width at half the maximum (FWHM) of the highest peak in the coefficient of variance histogram to be GUVs. Once selected, we collected the diameters and the counts of all of the GUVs from the tilescan.

### On surface imaging

We used a 20× Plan-Apochromat objective with a numerical aperture of 1.0 to observe the merging of close-packed buds. We captured 150 µm × 150 µm images ∼ 10 μm from the surfaces at an interval of 3.5 seconds for 7 minutes. We imaged two separate locations at 3 minutes and at 53 minutes on three independent samples. The pixel resolution was 0.119 µm and the confocal slice thickness was 1.5 μm. For the time-lapse images of close-packed buds on the surfaces we used a 10× Plan-Apochromat objective with a numerical aperture of 0.45. We collected 15 z-slices starting at the surface of the substrate and moving up in 2.8 μm increments. We collected z-stacks at 5 minutes, 30 minutes, 60 minutes, and 120 minutes. The area of the images was 425 µm × 425 µm, the pixel resolution was 0.265 µm, and the pinhole was set to 0.85 A.U. which resulted in a slice thickness of 5.6 μm. To obtain a 2-dimensional projection of the buds, we summed the z-slices using in ImageJ. For time lapse images of sparse buds, we used a 10× Plan-Apochromat objective with a numerical aperture of 0.45. The images covered an area of 340 µm × 340 µm with a pixel resolution of 0.265 µm, and the pinhole was set to 1 A.U. resulting in a slice thickness of 5.8 µm. We collected 11 slices per z-stack with an interval between stacks of 1 minute. We summed the slices of the z-stacks in ImageJ to create a 2-dimesional projection of the buds at each time point.

### Calculation of the average rate of merging

We used the point selection tool in ImageJ to mark and count the instances where two buds could clearly be seen merging to form a large bud. We estimate the average rate of merging by dividing the total number of merging events per unit area by the total time of observation.

### Analysis of incorporation of lipids

We measured the area of the buds using the point selection tool in ImageJ. For non-merging GUV buds, we traced the area of the bud at 5-minute intervals to obtain the diameter vs time. For buds that show an increase in diameter and merging, we traced the area of the final merged bud from the last frame. We then traced backwards in time toward the first frame to allow for the assignment of bud identities. We labeled the final merged bud as Bud 1. The three buds that merged to form Bud 1 were labeled Bud1a, Bud1b and Bud1c. Bud1b formed when three buds, Bud1ba, Bud1bb and Bud1bc merged.

## Results

### The distribution of GUV sizes is broad and increases with time

Figure 1 shows representative images of the GUVs harvested from each technique after 1 minute, 10 minutes, and 120 minutes. Qualitatively, the number of GUVs larger than 10 µm in diameter appear to increase with time for PAPYRUS and electroformation. Small GUVs between 1 µm and 10 µm are always present. Finding GUVs greater than 50 µm in diameter in typical fields of view is more common after 120 minutes of incubation for PAPYRUS and electroformation while finding GUVs larger than 50 µm in diameter is rare for gentle hydration on glass even for samples allowed to incubate for 120 minutes. It is interesting to note that there are significant numbers of GUVs for all three techniques even at the 1-minute stopping time.

**Figure 1.**
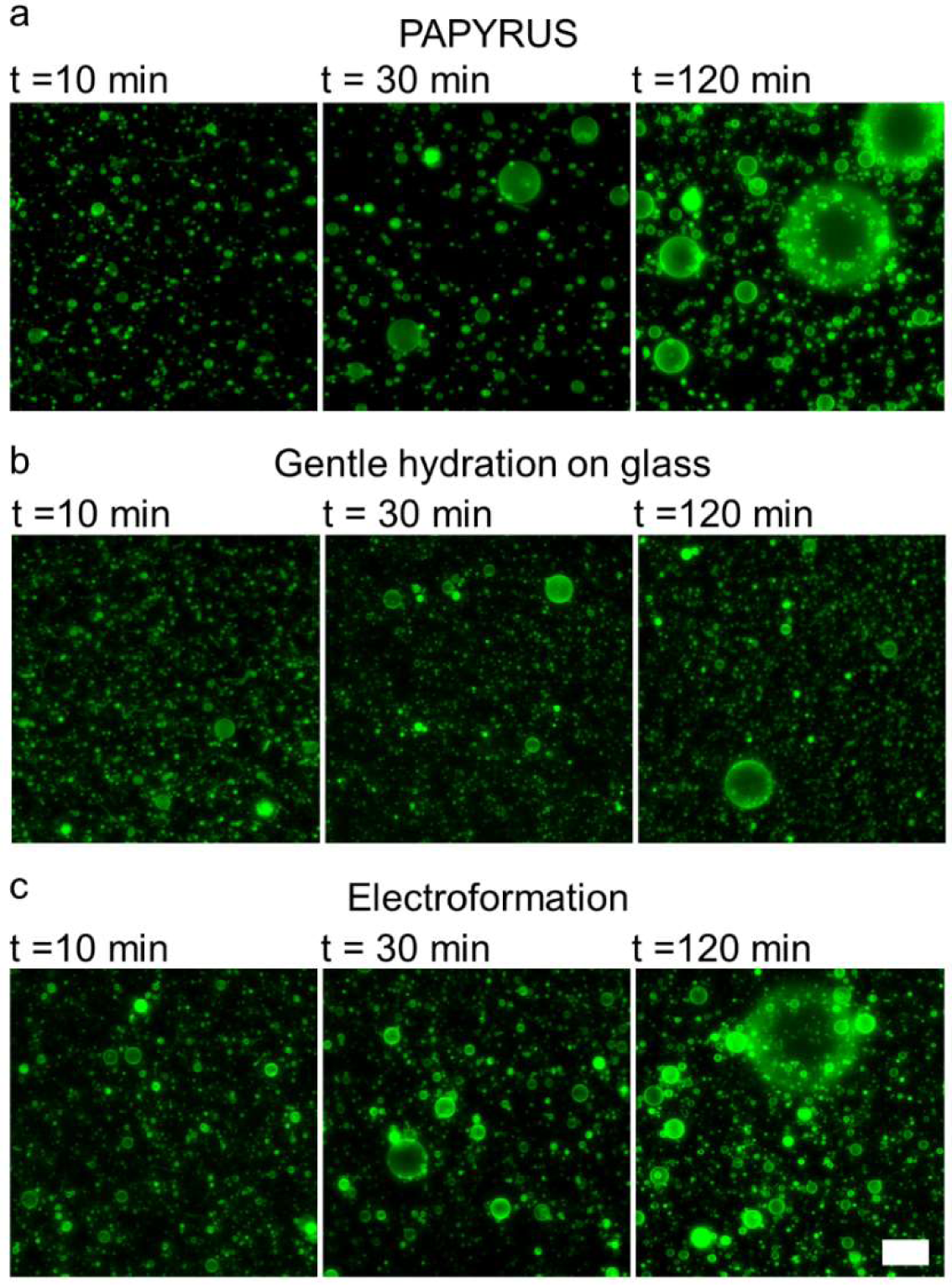
GUVs with large diameters are more abundant at longer incubation times. Representative confocal images of GUVs harvested from (a) PAPYRUS, (b) gentle hydration on glass, (c) electroformation. Scale bar 50 *μ*m.

Figure 2 shows histograms of the distribution of diameters of the GUVs for an incubation time of 1 minute, 10 minutes, and 120 minutes. The bin width is 1 µm and each bin is an average of three independent samples for each time point. The data is normalized by the amount of lipid mass, 10 µg, deposited on the substrates. The histograms for the other time points are shown in Figure S1, S2, and S3. Summary statistics such as the median diameter and variance are shown in Table S1,S2, and S3.

**Figure 2.**
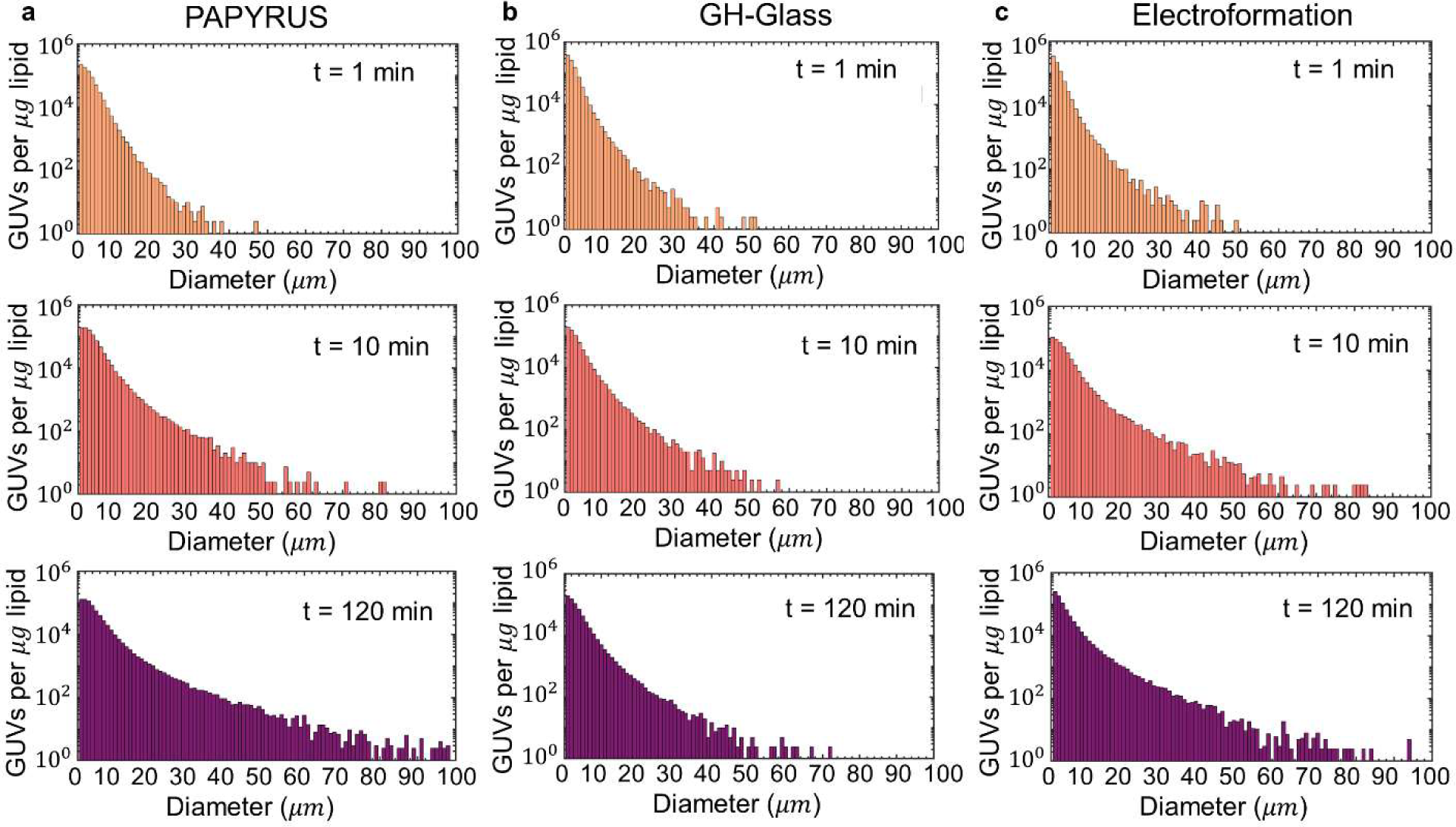
Distribution of diameters. The counts are normalized per *μ*g of lipid deposited for each sample. (a) PAPYRUS, (b) gentle hydration on glass, (c) electroformation. The stopping time is shown in the inset. The bin widths are 1 µm. Each bar represents an average from 3 independent samples.

All the samples show strikingly asymmetric distributions of diameters for all time points. The asymmetric distribution of diameters is a common feature of GUVs obtained through thin film hydration techniques such as from fabric,^26^ filter paper,^27^ glass,^13, 20, 28^ and gel-assisted hydration.^19^ In all samples and for all times, GUVs of small diameters are more abundant than GUVs of large diameters. There is no characteristic diameter of GUVs. We find that the distributions cannot be fit with common distributions used to describe dispersed particles that arise from classical nucleation and growth,^29^ coarsening,^30^ coalescence and fragmentation,^31^ nor those use to describe the evolution of nanoscale liposomes in bulk solution.^32^

To obtain further insight into the evolution of the GUVs with time, we divided our data into population classes based on diameter. Small GUVs (S-GUVs) have diameters between 1 *μm* ≤ *d* < 10 *μm* (Fig 3a-c), large GUVs (L-GUVs) have diameters between 10 *μm* ≤ *d* < 50 *μm* (Fig. 3d-f), and very large GUVs (VL-GUVs) have diameters greater than 50 *μ*m (Fig. 3g-i). The total number of GUVs is shown in (Fig. 3j-l). We find that the evolution of the total count of GUVs with time is dominated by the evolution of the counts of S-GUVs due to their large abundance relative to L-GUVs and VL-GUVs. The counts for S-GUVs for PAPYRUS and electroformation show non-monotonic behavior. The counts of S-GUVs decrease with time on glass. S-GUVs are more abundant for electroformation than the two gentle hydration techniques.

**Figure 3.**
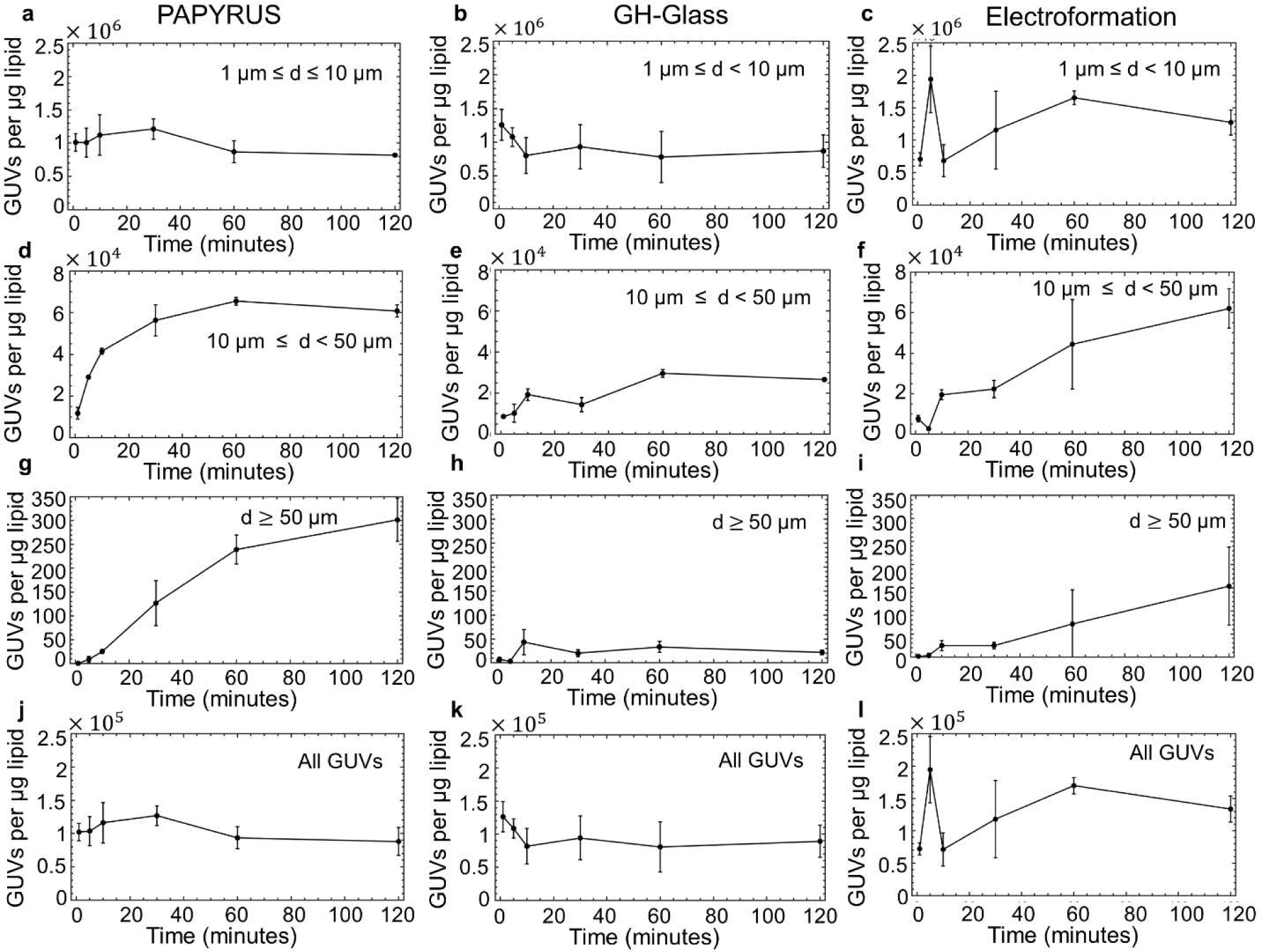
Evolution of GUV counts with time. (a-c) The number of small GUVs, (d-f) the number of large GUVs, (g-i) the number of very large GUVs and (j-l) the total number of GUVs. Each data point is a mean of three independent experiments and the error bars are the one standard deviation from the mean.

Unlike the evolution of S-GUVs, the number of L-GUVs, and VL-GUVs showed largely monotonic increases with time that reached a plateau within 2 hours for PAPYRUS and electroformation. Both these techniques had a larger number of VL-GUVs at 120 minutes compared to gentle hydration. PAPYRUS had more VL-GUVs compared to electroformation and gentle hydration.

To summarize, our data shows that obtaining the maximal number of large and very large GUVs requires 2 hours of incubation for electroformation and PAPYRUS. Obtaining small GUVs does not require long incubation times. Extending the incubation time to 3 hours does not change these results (Fig. S4, S5). The optimal incubation time to obtain S-GUVs is 30 minutes for PAPYRUS, 1 minute for gentle hydration on glass, and 5 minutes for electroformation. Obtaining a substantial number of VL-GUVs with gentle hydration does not appear to be possible. These results are useful for optimizing the incubation time to obtain GUVs of a desired diameter and places an upper limit on the diameter of GUVs that can be obtained.

### GUV yields plateau within one hour, coarsening can proceed for two hours

We next consider the evolution of the molar yield of GUVs with time. The molar yield measures the moles of lipid in the membranes of the population of harvested GUVs relative to the moles of lipids that were initially deposited on the surface.^20^ An increase in the molar yield indicates that lipid from the films on the surfaces have incorporated into the pool of buds that are harvested as GUVs. Drawing an analogy to chemical synthesis, the amount of lipids per unit area deposited onto the surface is the concentration of the ‘reactant’ and the amount of lipids in the membranes of the harvested GUVs is the ‘product’. We plot a stacked area plot of the molar yield versus time. The areas represent the portion of the yield that is comprised of S-GUVs, dark blue, L-GUVs, light blue, and VL-GUVs, white (Figure 4a-c). We note that although vesicles larger than 10 µm comprise less than 10 % of the population on a per count basis at 120 minutes for all three techniques, they make up to ∼ 60% of the molar yield from PAPYRUS, ∼ 33 % of the molar yield from gentle hydration on glass, and ∼ 54% of the molar yield from electroformation.

**Figure 4.**
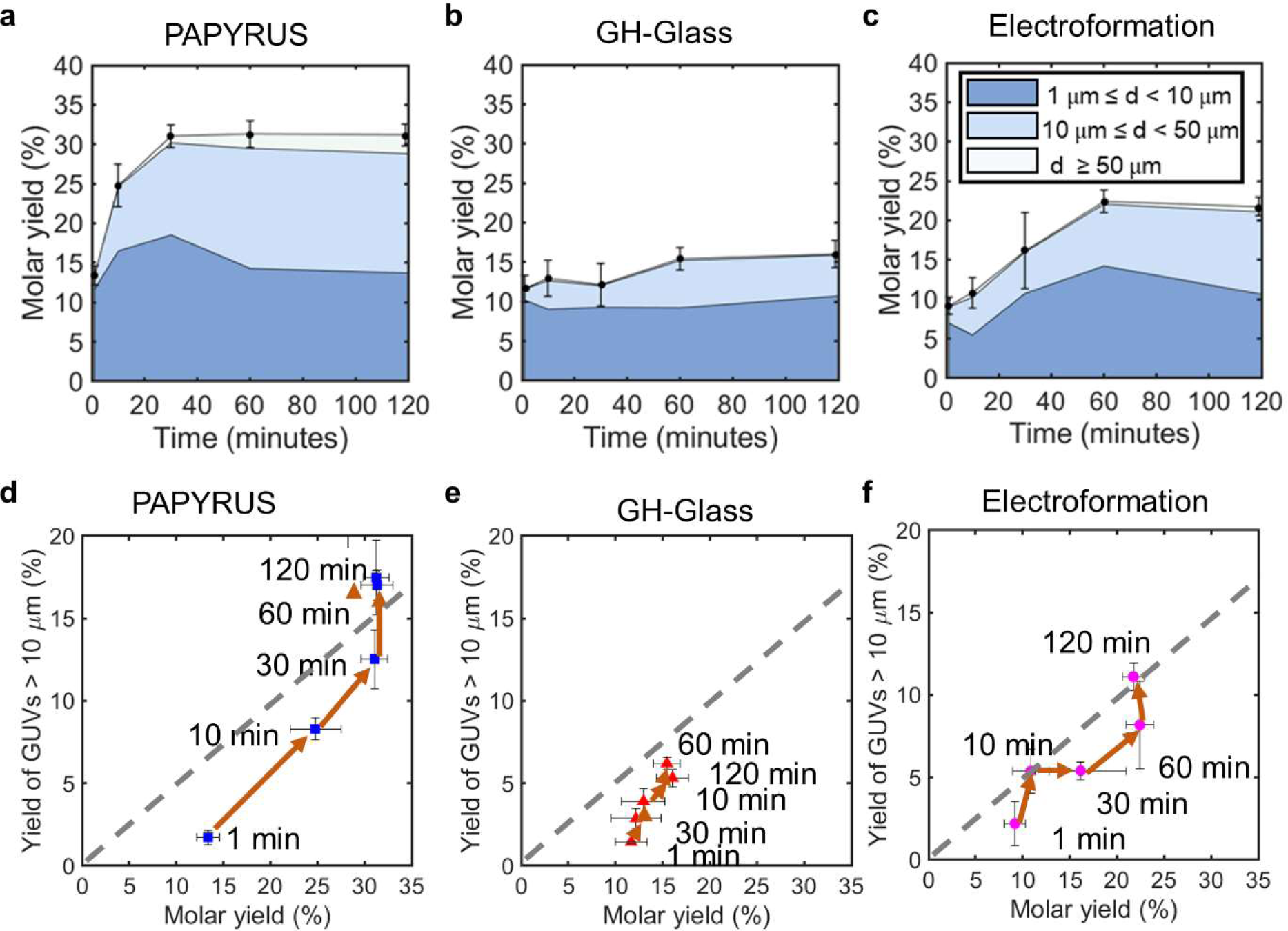
Evolution of molar yields with time. (a-c) Area plots showing the time evolution of the molar yield of GUVs. (a) PAPYRUS, (b) gentle hydration on glass, (c) electroformation. The areas show the percentage of the molar yield that is comprised from the different size classifications listed in the legend. (d-f) Scatter plots showing the molar yield from large GUVs (sizes greater than 10 *μ*m) versus the total molar yield. (d) PAPYRUS, (e) gentle hydration on glass, (f) electroformation. The orange arrows show progression of time. (a-f) Each point is an average of three independent experiments. The error bars show one standard deviation from the mean.

The total molar yield of GUVs monotonically increases with time before reaching a plateau, characteristic of a sigmoidal curve, for all three techniques. We empirically define the time to reach the plateau as one timepoint before the maximal yield. At the plateau, no new lipid is incorporated from the lipid films into the GUV sized buds. The yield of GUVs from PAPYRUS plateaus at 30 minutes and electroformation plateaus at 60 minutes. The molar yield of the GUVs obtained from PAPYRUS increases from 13 ± 1 % at 1 minute to 31 ± 1 % at 30 minutes. The molar yield of the GUVs obtained from electroformation increases from 9 ± 1 % at 1 minute to 22 ± 1 % at 60 minutes. In principle, gentle hydration on glass plateaus at 120 minutes. However, unlike the other two techniques, the difference in yield between the initial time point and the plateau is only 4 %, from 12 ± 2 % at 1 minute to 16 ± 2 % at 120 minutes. The average rate of lipid incorporation into the pool of GUVs from the initial measurement to the plateau was 7.7 × 10^-11^ mol min^-1^ for PAPYRUS, 4.7 × 10^-12^ mol min^-1^ for gentle hydration on glass and 2.8 × 10^-11^ mol min^-1^ for electroformation.

After reaching a plateau in total yield, the proportion of small, large, and very large GUVs changes between 60 minutes and 120 minutes for both PAPYRUS and electroformation, a result that can be seen from the change in the width of the stacked areas. This change in proportion of vesicles of different sizes is consistent with the coarsening of the buds.

We plot the molar yield of large GUVs versus the total molar yield of GUVs to further illustrate graphically the differences between the techniques (Fig. 4d-f). The *x* and *y* error bars are the standard deviation from the mean. The gray dashed line corresponds to the values where half of the GUVs are larger than 10 µm. In these plots, incorporation of lipid from the film into the pool of GUV-sized buds results in movement in the positive direction parallel to the *x*-axis. Coarsening of the buds to form buds of larger diameter at a constant amount of lipid in the pool results in movement in the positive direction parallel to the *y*-axis. Diagonal movements on the plot show both lipid incorporation and coarsening. The orange lines with arrowheads trace the progression of time. In these plots, the length of the path traced by the arrows from the intial time point to the final time point reflects the amount of evolution of the population of GUV sized buds with respect to time.

PAPYRUS shows a period of both incorporation of lipid and coarsening for the first 30 minutes and a period of significant coarsening with minimal incorporation of lipid between 30 and 60 minutes. Gentle hydration shows both limited incorporation of lipid and limited coarsening as can be seen by the short path length of the arrows. The time evolution of incorporaton of lipid and coarsening traces a more complex path for electroformation compared to the other two techniques. Electroformation shows a period of both lipid incorporation and coarsening from 1 to 10 minutes and then again from 30 to 60 minutes. From 10 to 30 minutes lipids incorporate into the GUVs with minimal coarsening. GUVs coarsen with no incorporation of lipid from 60 to 120 minutes.

### Observations of bud dynamics shows that the rate of merging decreases with time

To determine the local processes that give rise to the time evolution of diameters and yields of GUVs, we observe the buds on the surfaces by capturing four dimensional (*x, y, z, t*) time-lapse confocal images. At the typical concentration of 17 nmol/cm^2^ of lipids on the surface (10 μg), micrometer scale buds are abundant and close-packed on the surfaces of nanocellulose paper and the ITO-coated slides. We show a representative image using a color-coded *x-y* and *x-z* reconstruction of the buds on nanocellulose paper after 60 minutes of incubation (Figure 5a). The buds formed size-stratified layers ∼ 150 – 200 μm in thickness on the surface of the nanocellulose paper. Buds 1-5 μm in diameter were closer to the surface and buds > 5 μm were located further away from the surface. We see approximately two layers of buds on the surface of the ITO-coated slides for electroformation. We typically see only one layer of buds on the surface of the glass for gentle hydration.

**Figure 5.**
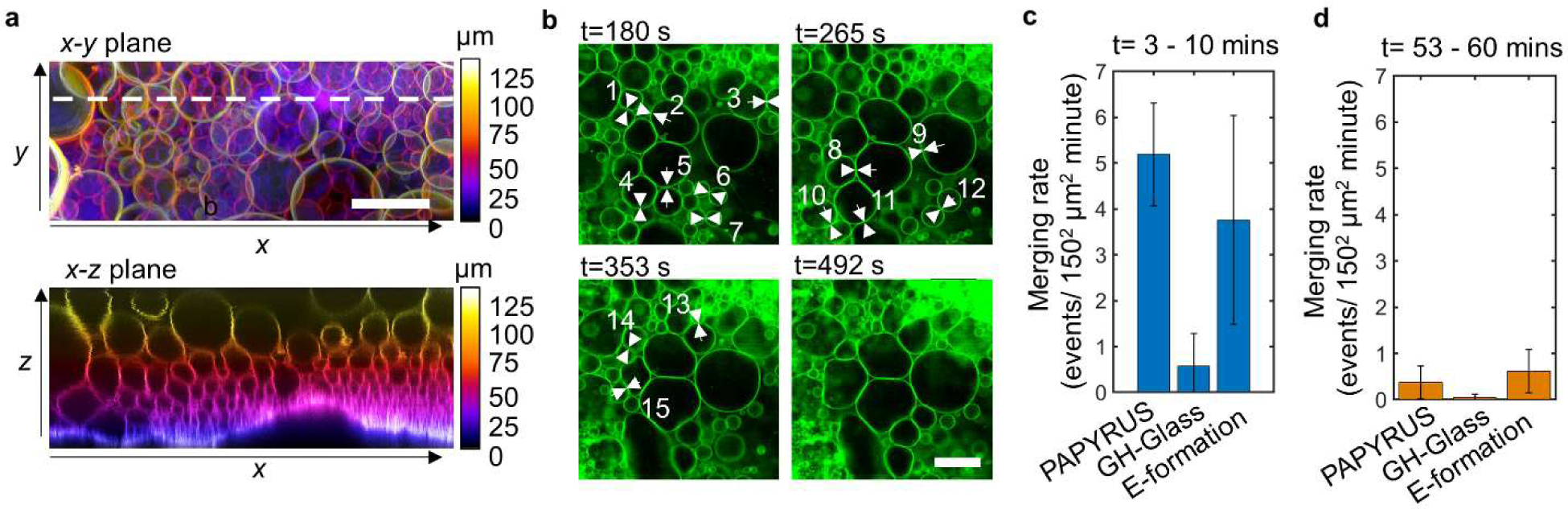
In situ analysis of close-packed buds on the surface. (a) Three-dimensional configuration of the buds on the surface. The upper panel shows a color-coded *x-y* projection using the sum slices method of the buds from a confocal z-stack. The lower panel shows an orthogonal *x-z* projection of the dashed line shown in the upper panel. The white layer at the bottom are high densities of small buds on the surface of paper. (b) Stills showing merging between buds. The white numbered arrowheads show distinct membranes that rearranged to merge. (c) The number of merging events between 3 to 10 minutes after hydration. (d) The number of merging events between 53 to 60 minutes after hydration. Each bar is the average from three different samples 150 μm × 150 μm in area and 10 μm from the surface of the substrates. The error bars are the standard deviation from the mean. Scale bar 20 μm.

At the typical concentration of 17 nmol/cm^2^ of lipids on the surface, the primary dynamical phenomena that we observe is merging of micrometer scale buds. Merging occurs when two adjacent buds, which are both connected to the lipid film, rearrange their membranes to form a single bud.^20^ The driving force for merging is the reduction in the total elastic energy of the connected buds on lipid film. At a constant total area, a film with fewer spherical buds of larger diameters has a lower total elastic energy than a film with more buds of smaller diameters.^20^ Additionally merging can be enhanced due to the action of an electric field.^24^ The buds remain connected to the lipid film on the surface until detached.^20, 33^

We probe the time dependence of the rate of merging by observing three regions in a 150 × 150 μm area approximately 10 μm above the surface of the paper between 3 and 10 minutes after hydration and between 53 and 60 minutes after hydration. A characteristic sequence of images is shown in Fig. 5b. Fig. 5c, d shows a bar plot of the number of merging events that occur over the observation time of 7 minutes. PAPYRUS had the highest number of merging events at 5 events per 22,500 μm^2^ per minute, followed by electroformation at 4 events per 22,500 μm^2^ per minute. Gentle hydration on glass had the lowest number of merging events at 0.5 events per 22,500 μm^2^ per minute. At 53 – 60 minutes (Fig. 5d), the rate of merging decreased to below one event per 22,500 μm^2^ per minute for all three techniques.

We also captured high-resolution time-lapse images of buds evolving on the surface (Fig. 6). Discerning the diameters, changes in the diameters, and observing the emergence of new buds was difficult due to the high density of buds. Further, these images favor large GUVs, while the more numerous small GUVs closer to the surface are obscured. This observation suggests care should be taken when extrapolating distributions obtained from two dimensional images of surfaces containing multiple layers of buds to the distribution of harvested GUVs.^22, 34–36^

**Figure 6.**
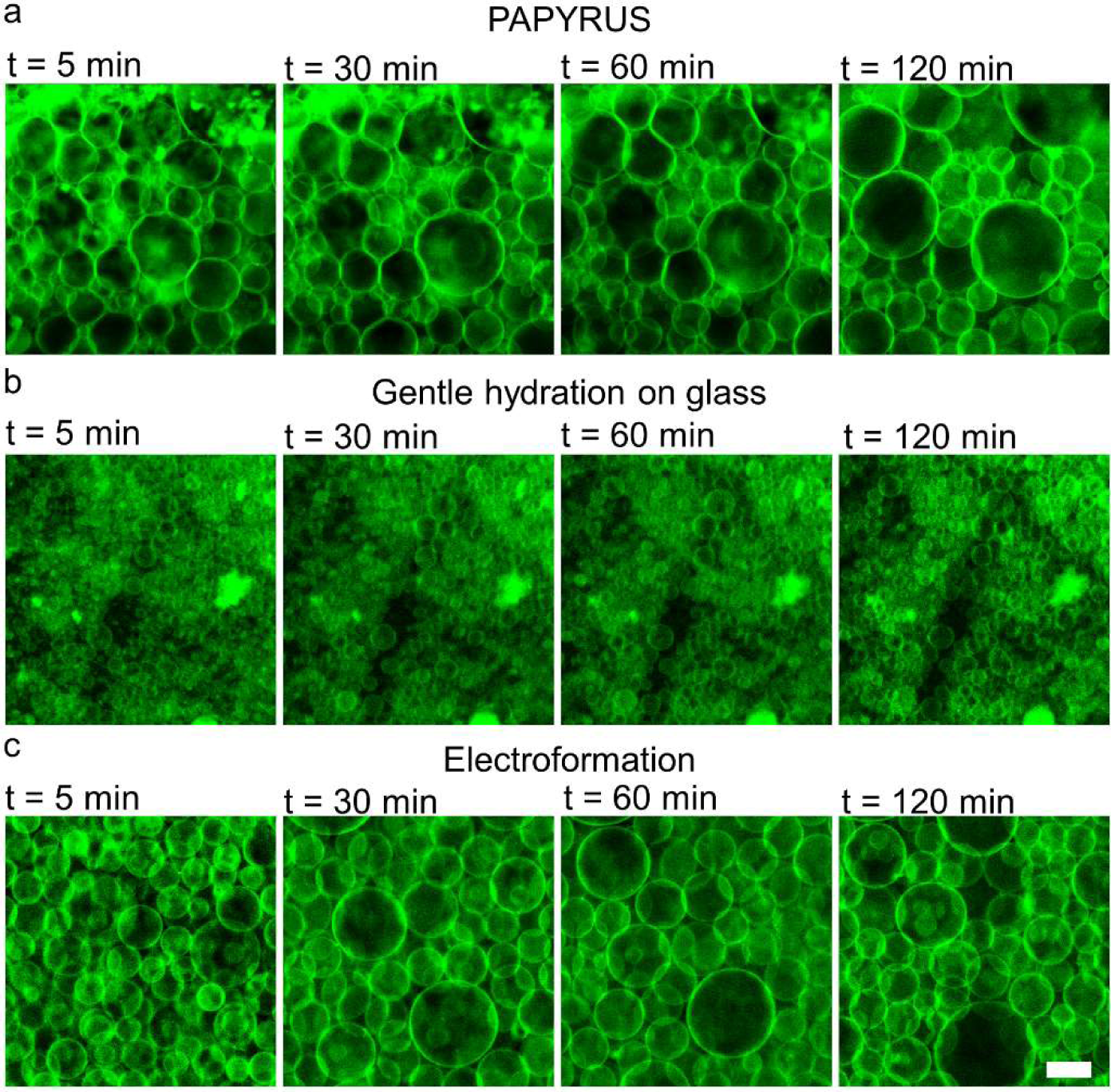
Confocal images showing the time evolution of buds. (a) PAPYRUS. (b) Gentle hydration on glass. (c) Electroformation. The images are sum projections of confocal z-stacks. Scale bar 25 *μ*m.

### Samples with sparse buds show lipid is incorporated into new and existing buds, merging is the primary mechanism to obtain large buds

We created surfaces with a sparse monolayer coverage of buds by depositing four times less lipid on the surface (4.25 nmol/cm^2^, 2.5 µg). We focused on the PAPYRUS technique since it produced the highest yield. We find that on these surfaces, two-dimensional projections can capture the dynamics of buds (Figure 7 a,b). Buds clearly appear larger after 60 minutes of incubation. Along with merging, we could now discern two other dynamical processes on the surface - i) emergence of new micrometer sized buds and ii) incorporation of lipid into existing buds resulting in an increase in the diameter of the buds.

**Figure 7.**
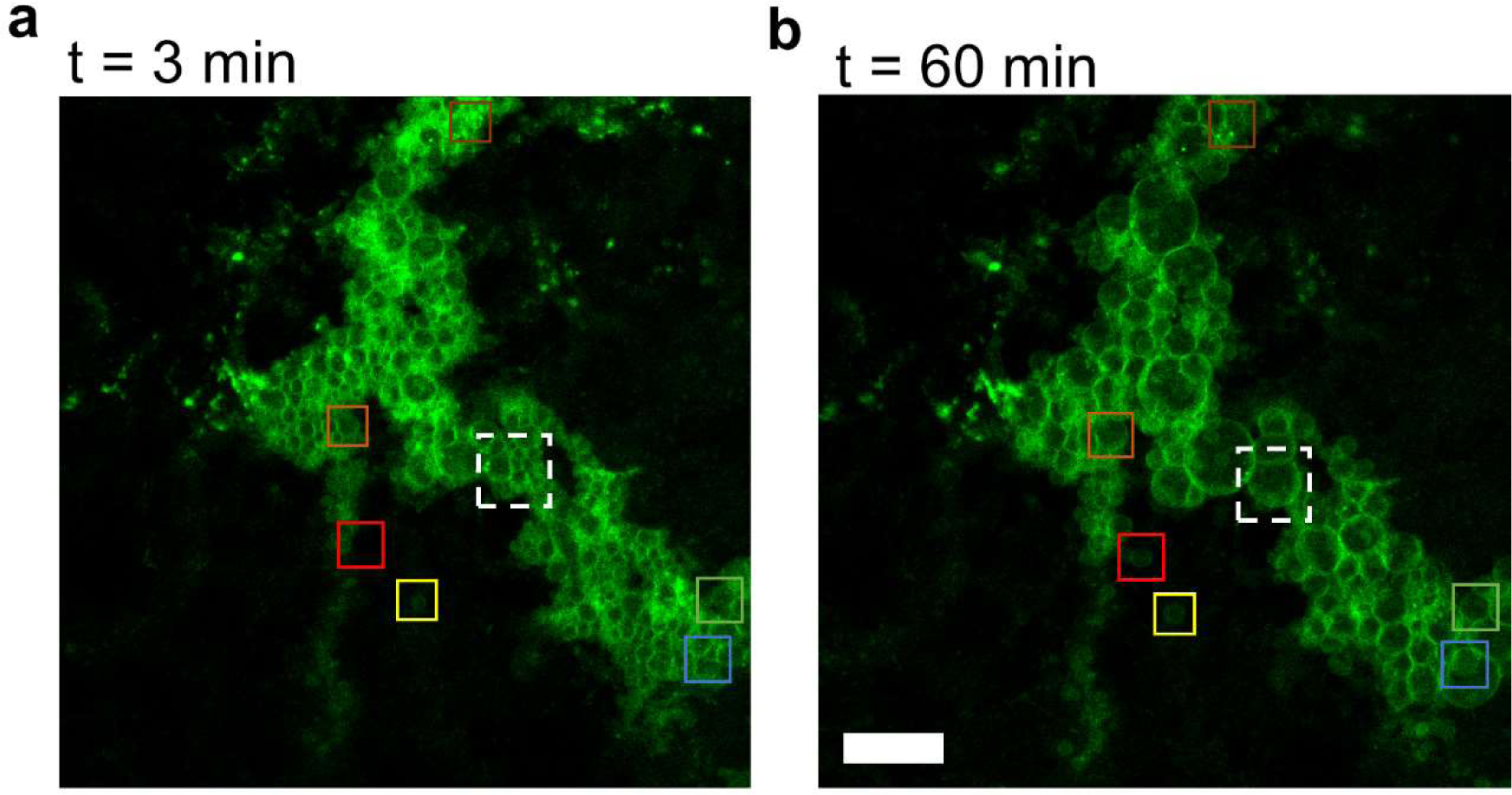
Stills of buds on paper surfaces with low surface concentration of lipids. The colored boxes correspond to the buds analyzed in Fig. 8a, the white dashed box corresponds to the cluster of buds analyzed in Fig 8b-d. Scale bar 50 µm.

We show examples of diameter versus time curves of six individual buds that do not appear to merge with their neighbors in Figure 8a. The locations of these buds are shown as colored boxes in Fig. 7. Bud 1, 2, 4, and 5 were already micrometer in diameter at the earliest time of observation. Bud 3 and 6 emerged 15 minutes after we began our observation. The 6 curves show a sigmoidal shape (Fig. 7d). Buds separated in location on the surface show different rates of lipid incorporation. The fastest rate of lipid incorporation was for Bud 3 at 9.7 × 10^-17^ mol min^-1^ and the slowest rate of incorporation was for Bud 1 at 1.9 × 10^-17^ mol min^-1^. On average the buds increase in diameter by 44 %. Buds moved from being classified as S-GUVs to L-GUVs. However, none of the buds had diameters that exceeded 20 µm.

**Figure 8.**
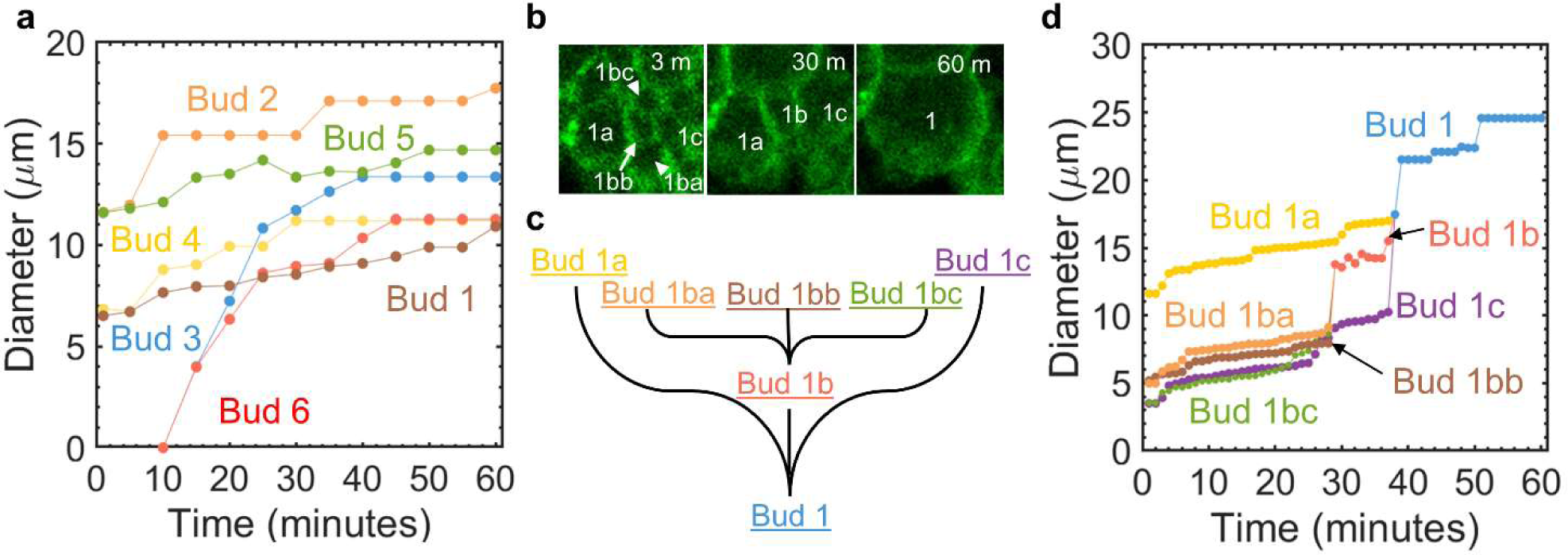
Incorporation of lipids into new and existing buds. (a) Diameter vs time trajectories for the 6 buds shown in Fig.7 that show no apparent micrometer scale merging. (b) Zoomed stills of the cluster of buds shown in Fig. 7. (c) Diagram of merging hierarchy. At the first time point, there are 5 buds. Bud 1ba, Bud1bb, Bud1bc merge to form Bud1b. Bud1a, Bud1b, and Bud1c merge to form Bud 1. (d) Diameter vs time trajectories of the buds showing that the buds increase in diameter. Merging results in step increase in bud diameters.

Indeed, even at this concentration of lipids, buds that increase in diameter without micrometer scale merging are rare. We find that most buds increase in diameter and then merge with their neighbors. Figure 8b shows zoomed stills of the evolution of 5 buds shown in the white dashed box in Figure 7. Figure 8c shows the hierarchy of merging. At the initial observation time, 5buds were present. Then Bud 1ba, Bud1bb, and Bud1bc merge at 29 minutes to form Bud 1b. Bud 1a, Bud1b, and Bud 1c merge at 38 minutes to form Bud 1. All the buds clearly show an increase in their diameter due to incorporation of new lipids punctuated by discontinuous step increases in diameter at merging events (Fig. 8d). Unlike buds separated in location, the rate of increase in diameter for these buds that are in close proximity is similar to each other. A combination of merging and incorporation of new lipids into micrometer scale buds allows the formation of GUVs with diameters greater than 25 µm.

To summarize, the dynamics of the buds on the surfaces, though complex, qualitatively match the observation of the evolution of the population of GUVs. Both the rate of merging of micrometer scale buds, which contributes to bud coarsening, and the rate of incorporation of new lipids into the buds, which contributes to the increase in yield, decrease with time. The equal importance of bud emergence, bud growth, and bud merging likely explains the complex shape of the distributions of diameters of GUVs.

### The budding and merging model explains the time evolution of GUVs

The underlying dynamics of the buds are clearly complex. Nevertheless, fundamental observation unites the thin film hydration techniques. Despite the different surface geometries and input of energy, all three techniques have relatively high initial yields that increase at different rates and then plateaus. We rationalize these results using the budding and merging model for the assembly of GUVs. In this model, GUV sized buds assemble from connected spherical buds that merge to form large buds on the thin lipid films.^20^ In this minimal thermodynamic model, the formation of spherical buds from lipid films depends on the geometry of the film and the size of the buds.^20^ Merging of the buds to form few buds of large sizes is always energetically favorable. The free energy change for the formation of spherical buds from flat bilayers is several hundreds to a thousand times that of ambient thermal energy.^20^ Small buds down to the nanoscale are expected to be more abundant since the energy to form a bud increases with the radius of the bud.^20^ Since there is input of energy above ambient during the moment of hydration due to heats of hydration and hydrodynamic flows, we expect buds of various sizes to form on the thin lipid films regardless of the geometry of the films. The substantial number of GUVs obtained within the relatively short incubation time of 1 minute after hydration for all three techniques is consistent with this expectation.

Beyond this initial time, the time evolution of yields diverges for the three techniques. For PAPYRUS, the bilayers are templated on the nanoscale cylindrical fibers of nanocellulose. The change in free energy is negative for the formation of nanoscale spherical buds from nanoscale cylindrical bilayers.^20^ Thus, the formation of nanoscale spherical buds is spontaneous compared to the ambient thermal energy scale.^20^ We suggest that the process of emergence of micrometer scale buds is explained when sufficient numbers of nanoscale buds merge with each other to form micrometer scale buds. The increase in diameter of the buds is consistent with nanoscale buds merging with already visible micrometer scale buds. Merging of the nanoscale buds explains our observed incorporation of new lipids into the pool of micrometer scale buds without requiring any further input of energy, resulting in an increase in GUV yield by an additional 20 % over the course of 120 minutes. The plateau that we observe likely reflects a depletion of the available pool of nanoscale buds.

In contrast, gentle hydration on glass shows both limited evolution of yields and limited coarsening of the buds compared to PAPYRUS. Gentle hydration on glass occurs in quiescent solution and does not have any obvious sources of energy. Since the formation of additional buds after the moment of hydration from flat bilayers is energetically costly, the limited increase in the yield of GUVs after the initial hydration is rationalized. For electroformation, the electric field inputs energy by acting on the charges in the lipid headgroup and solution.^14, 36–39^ The formation of additional buds due to the action of the electric field leads to the observed increase in yield of GUVs by an additional 11 % over the course of 120 minutes. We speculate the input of energy from the electric field causes the more complex time evolution of the buds (Fig. 4f) compared to PAPYRUS and gentle hydration (Fig.4d,e).

## Discussion

Quantitative experiments have unveiled significant insights into the assembly dynamics of GUVs using three distinct techniques: PAPYRUS, gentle hydration on glass, and electroformation. The time evolution of the molar yield for all three techniques follows sigmoidal curves, indicating a characteristic pattern of assembly. Furthermore, we have determined critical time thresholds beyond which no new lipid incorporates into GUV buds: 30 minutes for PAPYRUS, 120 minutes for gentle hydration, and 60 minutes for electroformation.

Surface imaging studies have provided compelling evidence that bud merging serves as the predominant mechanism for the generation of GUVs larger than 10 μm. Surprisingly, even at the earliest incubation time of 1 minute, numerous GUV buds are already present. However, achieving large GUVs necessitates longer incubation times of 1 to 2 hours for PAPYRUS and electroformation. Notably, gentle hydration has proven to be ineffective in obtaining large GUVs.

These findings open exciting prospects for further quantitative experimental and theoretical investigations into the mechanisms of self-assembly of GUVs. Furthermore, the newfound understanding of GUV assembly dynamics provides a pathway to tailor the production of giant vesicles with specific sizes for a range of applications. Thus, this research sets the stage for advancing the field of scalable GUV self-assembly and its practical implications.

## Supporting Information

Figure showing histograms of GUV diameters from different substrates at different times. Table showing summary statistics of the histograms. Figure S1-S5, Table S1-S3 (PDF).

## AUTHOR INFORMATION

### Author contributions

A.B.S and J.P conceived the study. J.P. performed experiments and analyzed the data. A.B.S. and J.P interpreted the data. J.P. prepared the figures and wrote the first draft of the manuscript. A.B.S wrote the final draft of the manuscript. All authors have given approval to the final version of the manuscript.

## Acknowledgements

This work was funded by the National Science Foundation through NSF CAREER DMR-1848573. The data in this work was collected, in part, with a confocal microscope acquired through the National Science Foundation MRI Award Number DMR-1625733.

## Conflict of Interest

The authors declare no conflict of interest.

## Supporting Information

### Supporting Figures

**Figure S1.**
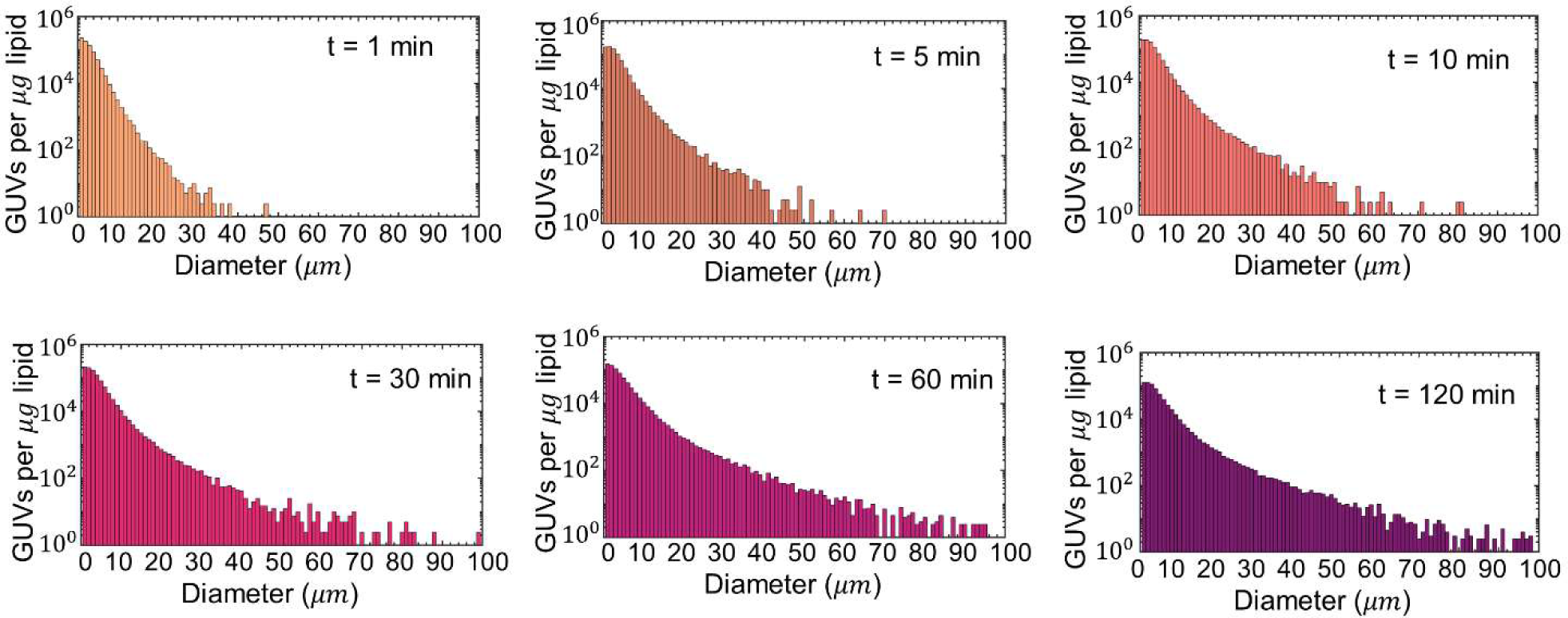
Histograms of GUV diameters for different incubation time points for PAPYRUS. Each histogram is the average of 3 independent repeats per time point. Note the logarithmic scale on the y-axis. Bin widths are 1 µm.

**Figure S2.**
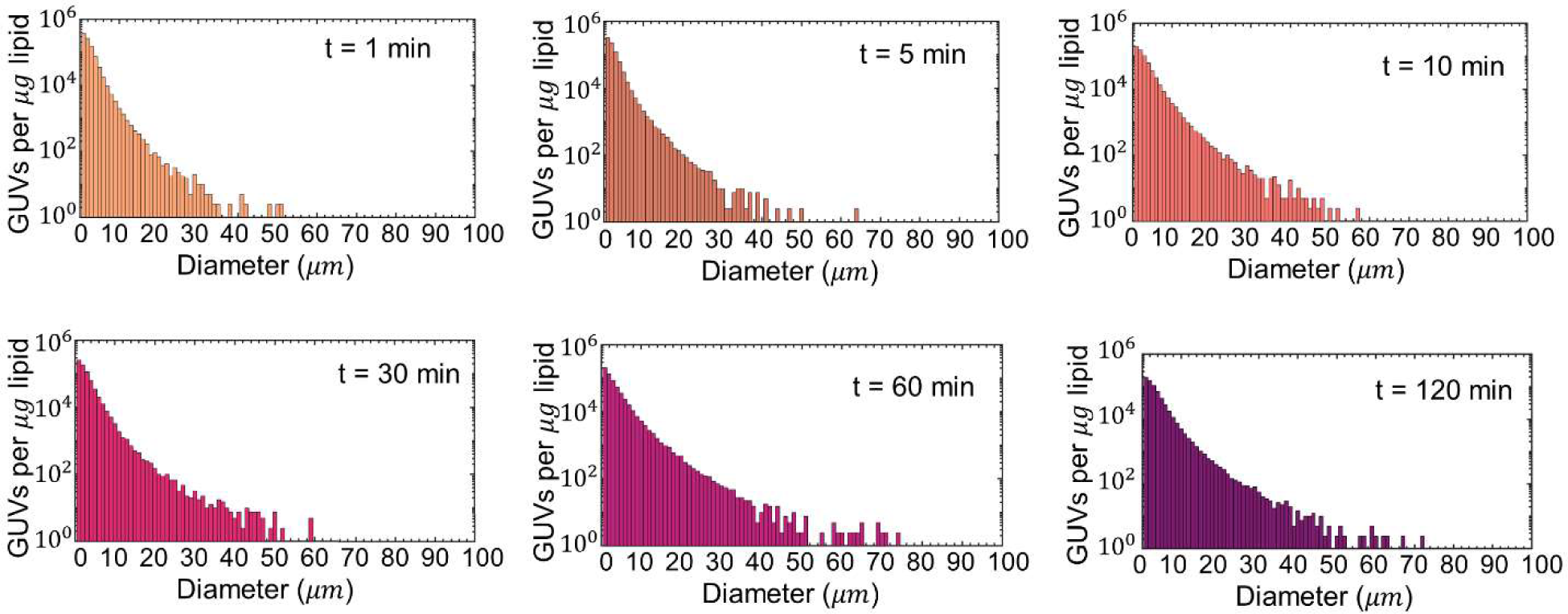
Histograms of GUV diameters for different incubation time points for gentle hydration on glass. Each histogram is the average of 3 independent repeats per time point. Note the logarithmic scale on the y-axis. Bin widths are 1 µm.

**Figure S3.**
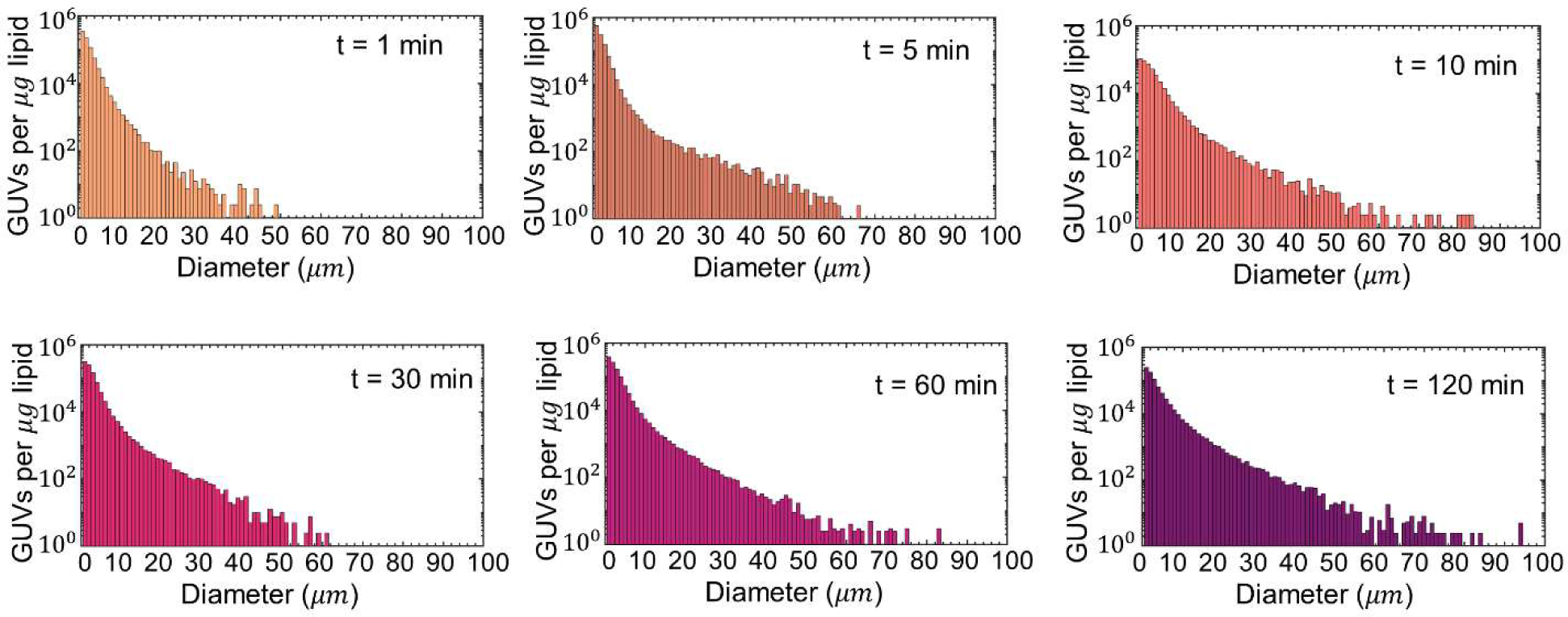
Histograms of GUV diameters for different incubation time points for electroformation. Each histogram is the average of 3 independent repeats per time point. Note the logarithmic scale on the y-axis. Bin widths are 1 µm.

**Figure S4.**
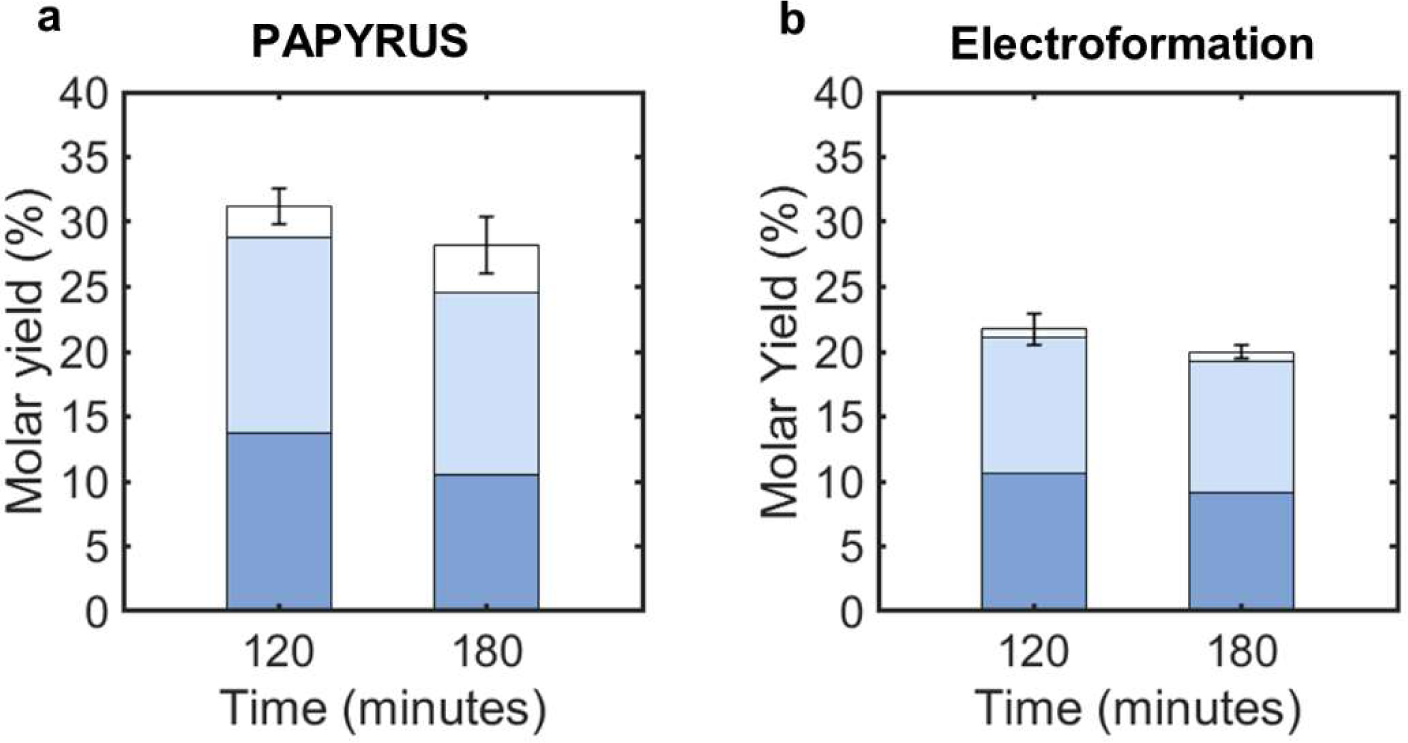
The molar yield after 180 minutes of incubation for (a) PAPYRUS and (b) electroformation. The data from 120 minutes is reproduced for comparison.

**Figure S5.**
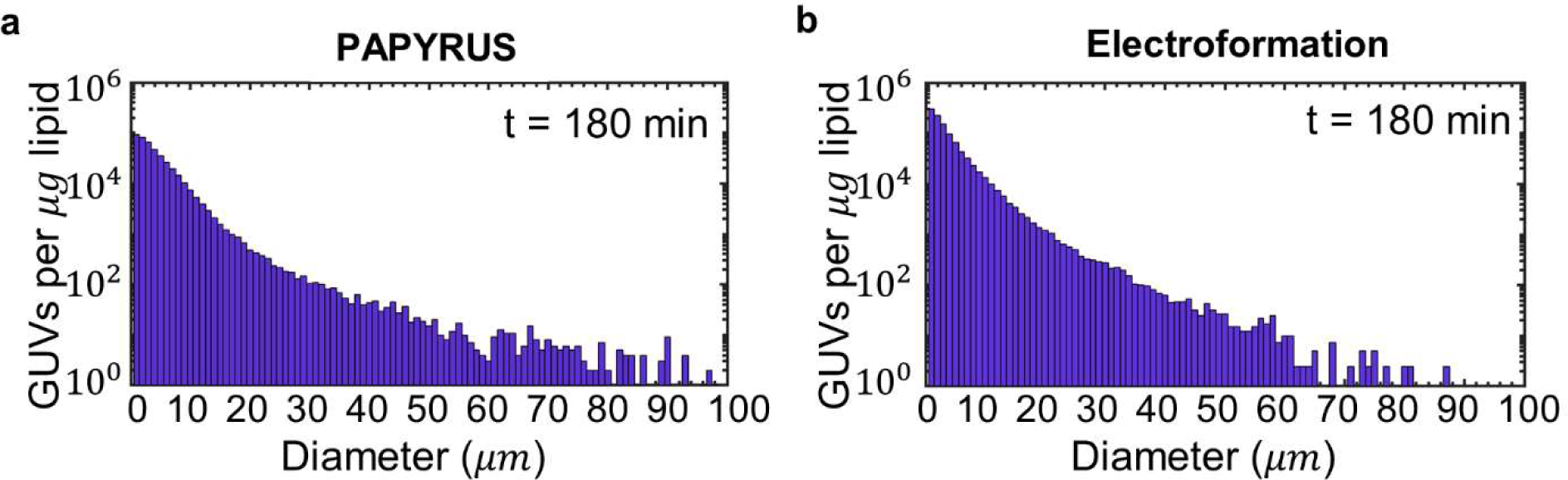
The histogram of diameters after 180 minutes of incubation for (a) PAPYRUS and (b) electroformation.

**Table S1.** Median diameter and variance of the GUVs for different incubation time points for PAPYRUS. Each value is an average of three independent repeats per time point. The errors are standard deviation from the mean.

| Time (minutes) | Median ($\mu m$) | Variance ($\mu m^2$) |
| --- | --- | --- |
| 1 | $2.8 \pm 0.3$ | $4.0 \pm 0.5$ |
| 5 | $3.3 \pm 0.2$ | $7 \pm 1$ |
| 10 | $3.3 \pm 0.2$ | $9 \pm 2$ |
| 30 | $3.4 \pm 0.3$ | $11 \pm 2$ |
| 60 | $3.6 \pm 0.4$ | $17 \pm 2$ |
| 120 | $3.6 \pm 0.2$ | $20 \pm 6$ |

**Table S2.** Median diameter and variance of the GUVs for different incubation time points for gentle hydration on glass. Each value is an average of three independent repeats per time point. The errors are standard deviation from the mean.

| Time (minutes) | Median ($\mu m$) | Variance ($\mu m^2$) |
| --- | --- | --- |
| 1 | $2.3 \pm 0.1$ | $3.1 \pm 0.4$ |
| 5 | $2.3 \pm 0.1$ | $3.4 \pm 0.5$ |
| 10 | $2.7 \pm 0.1$ | $8 \pm 3$ |
| 30 | $2.5 \pm 0.1$ | $6 \pm 1$ |
| 60 | $3.0 \pm 0.5$ | $10 \pm 3$ |
| 120 | $2.9 \pm 0.3$ | $8 \pm 2$ |

**Table S3.** Median diameter and variance of the GUVs for different incubation time points for electroformation. Each value is an average of three independent repeats per time point. The errors are standard deviation from the mean.

| Time (minutes) | Median ($\mu m$) | Variance ($\mu m^2$) |
| --- | --- | --- |
| 1 | $2.6 \pm 0.1$ | $3 \pm 1$ |
| 5 | $2.0 \pm 0.1$ | $2 \pm 1$ |
| 10 | $2.6 \pm 0.3$ | $10 \pm 1$ |
| 30 | $2.5 \pm 0.1$ | $7 \pm 2$ |
| 60 | $2.5 \pm 0.1$ | $7 \pm 3$ |
| 120 | $2.8 \pm 0.1$ | $13 \pm 1$ |

## Notes

### Competing Interest Statement

The authors have declared no competing interest.

## References

1. Dolder, N., Müller, P. & von Ballmoos, C. Experimental platform for the functional investigation of membrane proteins in giant unilamellar vesicles. Soft Matter 18, 5877– 5893 (2022).

2. Steinkühler, J. et al. Controlled division of cell-sized vesicles by low densities of membrane-bound proteins. Nat. Commun. 11, 1–11 (2020).

3. Xin, W., Wu, H., Grason, G. M. & Santore, M. M. Switchable positioning of plate-like inclusions in lipid membranes : Elastically mediated interactions of planar colloids in 2D fluids. Sci. Adv. 8, eabf1943 (2021).

4. Gözen, I. et al. Protocells: Milestones and Recent Advances. Small 2106624 (2022). doi:10.1002/smll.202106624

5. Cho, E. & Lu, Y. Compartmentalizing cell-free systems: Toward creating life-like artificial cells and beyond. ACS Synth. Biol. 9, 2881–2901 (2020).

6. Wang, X., Du, H., Wang, Z., Mu, W. & Han, X. Versatile Phospholipid Assemblies for Functional Synthetic Cells and Artificial Tissues. Adv. Mater. 33, 1–23 (2021).

7. Lussier, F., Staufer, O., Platzman, I. & Spatz, J. P. Can Bottom-Up Synthetic Biology Generate Advanced Drug-Delivery Systems? Trends Biotechnol. 39, 445–459 (2021).

8. Teixeira, L. et al. Use of giant unilamellar lipid vesicles as antioxidant carriers in in vitro culture medium of bovine embryos. Sci. Rep. 1–12 (2022). doi:10.1038/s41598-022-14688-8

9. Perrier, D. L., Rems, L. & Boukany, P. E. Lipid vesicles in pulsed electric fields: Fundamental principles of the membrane response and its biomedical applications. Adv. Colloid Interface Sci. 249, 248–271 (2017).

10. Has, C. & Sunthar, P. A comprehensive review on recent preparation techniques of liposomes. J. Liposome Res. 1–30 (2019). doi:10.1080/08982104.2019.1668010

11. Nair, K. S. & Bajaj, H. Advances in giant unilamellar vesicle preparation techniques and applications. Adv. Colloid Interface Sci. 318, 102935 (2023).

12. Bangham, A. D., Standish, M. M. & Watkins, J. C. Diffusion of univalent ions across the lamellae of swollen phospholipids. J. Mol. Biol. 13, 238–252 (1965).

13. Reeves, J. P. & Dowben, R. M. Formation and properties of thin-walled phospholipid vesicles. J. Cell. Physiol. 73, 49–60 (1969).

14. Dimitrov, D. S. & Angelova, M. I. Lipid swelling and liposome formation mediated by electric fields. Bioelectrochemistry Bioenerg. 253, 323–336 (1988).

15. Pereno, V. et al. Electroformation of Giant Unilamellar Vesicles on Stainless Steel Electrodes. ACS Omega 2, 994–1002 (2017).

16. Angelova, M., Soleau, S. & Méléard, P. Preparation of giant vesicles by external AC electric fields. Kinetics and applications. Progr Colloid Polym Sci 89, 127–131 (1992).

17. Horger, K. S., Estes, D. J., Capone, R. & Mayer, M. Films of agarose enable rapid formation of giant liposomes in solutions of physiologic ionic strength. J. Am. Chem. Soc. 131, 1810–1819 (2009).

18. Weinberger, A. et al. Gel-assisted formation of giant unilamellar vesicles. Biophys. J. 105, 154–164 (2013).

19. Cooper, A., Girish, V. & Subramaniam, A. B. Osmotic Pressure Enables High-Yield Assembly of Giant Vesicles in Solutions of Physiological Ionic Strengths. Langmuir 39, 5579–5590 (2023).

20. Pazzi, J. & Subramaniam, A. B. Nanoscale Curvature Promotes High Yield Spontaneous Formation of Cell-Mimetic Giant Vesicles on Nanocellulose Paper. ACS Appl. Mater. Interfaces 12, 56549–56561 (2020).

21. Dimitrov, D. S. & Angelova, M. I. Lipid swelling and liposome formation on solid surfaces in external electric fields. Progr Colloid Polym. Sci 73, 48–56 (1987).

22. Politano, T. J., Froude, V. E., Jing, B. & Zhu, Y. AC-electric field dependent electroformation of giant lipid vesicles. Colloids Surfaces B Biointerfaces 79, 75–82 (2010).

23. Shimanouchi, T., Umakoshi, H. & Kuboi, R. Kinetic study on giant vesicle formation with electroformation method. Langmuir 25, 4835–4840 (2009).

24. Micheletto, Y. M. S., Marques, C. M., Silveira, N. P. Da & Schroder, A. P. Electroformation of Giant Unilamellar Vesicles: Investigating Vesicle Fusion versus Bulge Merging. Langmuir 32, 8123–8130 (2016).

25. Shimanouchi, T., Umakoshi, H. & Kuboi, R. Growth behavior of giant vesicles using the electroformation method: Effect of proteins on swelling and deformation. J. Colloid Interface Sci. 394, 269–276 (2013).

26. Girish, V., Pazzi, J., Li, A. & Subramaniam, A. B. Fabrics of Diverse Chemistries Promote the Formation of Giant Vesicles from Phospholipids and Amphiphilic Block Copolymers. Langmuir 35, 9264–9273 (2019).

27. Pazzi, J., Xu, M. & Subramaniam, A. B. A. B. Size distributions and yields of giant vesicles assembled on cellulose papers and cotton fabric. Langmuir 35, 7798–7804 (2018).

28. Karal, M. A. S. et al. Electrostatic interaction effects on the size distribution of self-assembled giant unilamellar vesicles. *Phys*. Rev. E 101, 1–11 (2020).

29. Thanh, N. T. K., Maclean, N. & Mahiddine, S. Mechanisms of Nucleation and Growth of Nanoparticles in Solution. Chem. Rev. 114, 7610–7630 (2014).

30. P.W. Voorhees. The Theory of Ostwald Ripening. J. Stat. Phys. 38, 231–252 (1985).

31. Spicer, P. T. & Sotiris E. Pratsinis. Coagulation and fragmentation: Universal steady-state particle-size distribution. AIChE J. 42, 1612–1620 (1996).

32. Huang, C., Quinn, D., Sadovsky, Y., Suresh, S. & Hsia, K. J. Formation and size distribution of self-assembled vesicles. Proc. Natl. Acad. Sci. U. S. A. 114, 2910–2915 (2017).

33. Estes, D. J. & Mayer, M. Giant liposomes in physiological buffer using electroformation in a flow chamber. Biochim. Biophys. Acta - Biomembr. 1712, 152–160 (2005).

34. Boban, Z., Mardešić, I., Subczynski, W. K. & Raguz, M. Giant unilamellar vesicle electroformation: What to use, what to avoid, and how to quantify the results. Membranes (Basel). 11, (2021).

35. Ghellab, S. E., Mu, W., Li, Q. & Han, X. Prediction of the size of electroformed giant unilamellar vesicle using response surface methodology. Biophys. Chem. 253, 106217 (2019).

36. Li, Q., Wang, X., Ma, S., Zhang, Y. & Han, X. Electroformation of giant unilamellar vesicles in saline solution. Colloids Surfaces B Biointerfaces 147, 368–375 (2016).

37. Seiwert, J. & Vlahovska, P. M. Instability of a fluctuating membrane driven by an ac electric field. *Phys. Rev. E - Stat. Nonlinear*, Soft Matter Phys. 87, 1–13 (2013).

38. Sens, P. & Isambert, H. Undulation instability of lipid membranes under an electric field. Phys. Rev. Lett. 88, 128102 (2002).

39. Dimova, R. et al. Giant vesicles in electric fields. Soft Matter 3, 817–827 (2007).

